# Exploration of the *Neisseria* resistome reveals resistance mechanisms in commensals that may be acquired by *N. gonorrhoeae* through horizontal gene transfer

**DOI:** 10.1101/2020.07.30.228593

**Authors:** Michael A. Fiore, Jordan C. Raisman, Narayan H. Wong, André O. Hudson, Crista B. Wadsworth

## Abstract

Non-pathogenic *Neisseria* have repeatedly been demonstrated to transfer antibiotic resistance genes to their pathogenic relative, *Neisseria gonorrhoeae.* However, the resistance genotypes and subsequent phenotypes of non-pathogens within the genus have been studied and described less frequently. Here, we use Etests to characterize the minimum inhibitory concentrations (MICs) of a panel of *Neisseria* (n=26) – including several commensal species – acquired from the CDC & FDA’s Antibiotic Resistance (AR) Isolate Bank to a suite of diverse antibiotics. We furthermore use whole genome sequencing and the Comprehensive Antibiotic Resistance Database (CARD) Resistance Gene Identifier (RGI) platform to predict possible causal resistance-encoding mutations. Within this panel, resistant isolates to all tested antimicrobials including penicillin (n=5/26), ceftriaxone (n=2/26), cefixime (n=3/26), tetracycline (n=10/26), azithromycin (n=11/26), and ciprofloxacin (n=4/26) were found. In total we identify 63 distinct mutations predicted by RGI to be involved in resistance. The presence of several of these mutations had clear associations with increases in MIC such as: DNA gyrase subunit A (*gyrA*) (S91F) and ciprofloxacin, tetracycline resistance protein (*tetM*) and 30S ribosomal protein S10 (*rpsJ*) (V57M) and tetracycline, and TEM-type β-lactamases and penicillin. However, mutations with strong associations to macrolide and cephalosporin resistance were not conclusive. This work serves as an initial exploration into the resistance-encoding mutations harbored by non-pathogenic *Neisseria,* which will ultimately aid in prospective surveillance for novel resistance mechanisms that may be rapidly acquired by *N. gonorrhoeae*.

## Introduction

Emergence of antibiotic resistance in pathogenic bacteria presents a challenge clinically for the successful treatment of infections, and is a global threat to public health. In the United States alone, antibiotic resistant bacteria cause an estimated 2.8 million infections and 35,000 deaths each year [1]. While resistance can arise in bacteria via *de novo* mutations, it can also be horizontally transferred from environmental [2–4], animal [5–7], or human-associated [8–10] microbial communities. Thus, profiling the resistome, or the collection of all of the antibiotic resistance mechanisms available to particular bacterial species [4], is an important step in prospective surveillance for novel resistance-encoding mutations that may be rapidly acquired by pathogens.

There are a number of mechanisms that are employed by bacteria to circumvent the efficacy of antibiotics. These include but are not limited to; reduced permeability and decreased drug influx, target modification, antibiotic degradation, and increased efflux through pumps [11]. The threat of rapid resistance acquisition via horizontal gene transfer (HGT) of all of these mechanisms is exponentially amplified in bacteria that are naturally competent and highly recombinogenic such as the *Neisseria*. The genus *Neisseria* is composed of several closely related Gram-negative species, which are typically isolated from the naso- and oropharynx of humans and animals. While most species are considered commensals or ‘accidental pathogens’, only one, *N. gonorrhoeae* is an obligate human pathogen which colonizes the additional sites of the urogenital tract and rectum, and causes the sexually transmitted infection gonorrhea. Concerningly, antimicrobial resistance is an increasing problem within *N. gonorrhoeae,* with over half of all of the 550,000 reported infections in the U.S. in 2017 resistant to at least one antibiotic [1], and treatment failures to the recommended azithromycin and ceftriaxone combination therapy reported internationally as of 2018 [12,13].

One of the likely reasons for the high prevalence of resistance in *N. gonorrhoeae* is due to its natural competence for transformation, preferentially with *Neisseria*-specific DNA, allowing for extensive intragenus gene exchange and quick access to new adaptive solutions [14]. Gene acquisition from commensal *Neisseria* is evidenced by widespread genomic mosaicism in *N. gonorrhoeae*, whereby specific mutations or haplotypes have been shown to have been inherited from its close relatives [15–19]. Furthermore, mutations encoding resistance to both azithromycin [10,20] and third-generation cephalosporins [21,22] have been demonstrated to have been transferred from non-pathogenic *Neisseria* to *N. gonorrhoeae*. Ultimately, these documented cases of widespread gene exchange between the *Neisseria* highlights the importance of surveying the non-pathogenic members of the genus for the resistance mechanisms that they harbor and may potentially share with their pathogen relative.

In this study, we set out to document both phenotypic and genotypic resistance across a panel of commensal *Neisseria* acquired from the CDC & FDA’s Antibiotic Resistance (AR) Isolate Bank – a bacterial strain collection resource for studying antibiotic resistance [23]. We phenotype this panel (n=26) to multiple classes of antimicrobials (beta-lactams, a macrolide, fluoroquinolone, and tetracycline), and report their minimum inhibitory concentrations (MIC). To link any observed increases in resistance to the possible underlying genetic contributors, we furthermore sequence the genomes of these isolates and use the Comprehensive Antibiotic Resistance Database (CARD) Resistance Gene Identifier (RGI) platform to predict possible causal resistance-encoding mutations.

## Results

### Characterization of phenotypic resistance in the AR Isolate Bank Neisseria panel

A total of 26 *Neisseria* isolates were obtained from the CDC & FDA’s Antibiotic Resistance (AR) Isolate Bank from the *Neisseria* species MALDI-TOF Verification panel (Table 1). Characterization of antimicrobial susceptibility to penicillin, ceftriaxone, cefixime, tetracycline, azithromycin, and ciprofloxacin was conducted using the Etest method (Table 1). For the gonococci within the panel (n=6), three were resistant to multiple antibiotic classes. *N. gonorrhoeae* AR Bank # 0936 was resistant to tetracycline, ciprofloxacin, and was just below the reduced susceptibility threshold for azithromycin (1.5 μg/mL with the breakpoint at 2 μg/mL); *N. gonorrhoeae* AR Bank # 0937 was resistant to penicillin, ciprofloxacin, and just below the reduced susceptibility threshold for tetracycline (1.5 μg/mL with the breakpoint at 2 μg/mL); and *N. gonorrhoeae* AR Bank # 0938 was resistant to penicillin, tetracycline, and ciprofloxacin.

**Table 1.** Minimum inhibitory concentrations of *Neisseria spps.* measured by Etest.

| AR Bank # | Species | Minimum inhibitory concentration (MIC)<sup>a,b</sup> |  |  |  |  |  |
| --- | --- | --- | --- | --- | --- | --- | --- |
|  |  | Penicillin (PEN) | Ceftriaxone (CRO) | Cefixime (CFX) | Tetracycline (TET) | Azithromycin (AZI) | Ciprofloxacin (CIP) |
| AR-0933 | *Neisseria gonorrhoeae* | 0.19 | 0.004 | < 0.016 | 1.5 | 0.25 | 0.004 |
| AR-0934 | *Neisseria gonorrhoeae* | 0.38 | 0.016 | 0.023 | 1.5 | 0.125 | 0.006 |
| AR-0935 | *Neisseria gonorrhoeae* | 0.19 | 0.006 | < 0.016 | 0.75 | 0.064 | 0.006 |
| AR-0936 | *Neisseria gonorrhoeae* | 0.38 | 0.047 | < 0.016 | **<u>2</u>** | 1.5 | **<u>≥ 32</u>** |
| AR-0937 | *Neisseria gonorrhoeae* | **<u>≥ 32</u>** | 0.016 | < 0.016 | 1.5 | 0.38 | **<u>6</u>** |
| AR-0938 | *Neisseria gonorrhoeae* | **<u>≥ 32</u>** | 0.008 | < 0.016 | **<u>32</u>** | 0.094 | **<u>4</u>** |
| AR-0943 | *Neisseria bacilliformis* | **<u>32</u>** | **<u>16</u>** | **<u>0.5</u>** | **<u>4</u>** | **<u>4</u>** | **<u>6</u>** |
| AR-0944 | *Neisseria cinerea* | 0.38 | 0.094 | < 0.016 | **<u>2</u>** | **<u>8</u>** | 0.032 |
| AR-0945 | *Neisseria elongata* | 0.25 | 0.094 | 0.032 | 0.38 | 0.5 | 0.25 |
| AR-0946 | *Neisseria lactamica* | 0.75 | 0.023 | 0.25 | 0.75 | 1.5 | 0.008 |
| AR-0947 | *Neisseria oralis* | 1.5 | 0.047 | 0.064 | 1.5 | 1.5 | 0.016 |
| AR-0948 | *Neisseria canis* | 0.25 | 0.008 | < 0.016 | 0.5 | 0.38 | 0.008 |
| AR-0949 | *Neisseria macacae* | 1.5 | 0.064 | 0.064 | **<u>2</u>** | **<u>8</u>** | 0.125 |
| AR-0950 | *Neisseria macacae* | **<u>3</u>** | 0.047 | 0.125 | **<u>3</u>** | **<u>8</u>** | 0.032 |
| AR-0951 | *Neisseria mucosa* | 0.25 | 0.032 | 0.125 | 1 | **<u>3</u>** | 0.023 |
| AR-0952 | *Neisseria macacae* | 0.38 | 0.023 | 0.047 | **<u>2</u>** | 0.5 | 0.012 |
| AR-0953 | *Neisseria subflava* | 1.5 | 0.019 | 0.38 | 0.5 | **<u>2</u>** | 0.75 |
| AR-0954 | *Neisseria subflava* | **<u>3</u>** | 0.25 | **<u>0.5</u>** | **<u>48</u>** | **<u>4</u>** | 0.047 |
| AR-0955 | *Neisseria subflava* | 1 | 0.032 | 0.064 | **<u>6</u>** | **<u>12</u>** | 0.125 |
| AR-0956 | *Neisseria subflava* | 1.5 | 0.125 | 0.25 | 1 | **<u>6</u>** | 0.75 |
| AR-0957 | *Neisseria subflava* | 1 | 0.047 | 0.064 | **<u>4</u>** | **<u>8</u>** | 0.064 |
| AR-0958 | *Neisseria weaveri* | 0.38 | 0.064 | 0.047 | 0.5 | 0.25 | 0.006 |
| AR-0959 | *Neisseria weaveri* | 1 | 0.19 | 0.023 | 0.75 | 0.75 | 0.023 |
| AR-0960 | *Kingella denitrificans* | 0.32 | 0.064 | 0.25 | 0.75 | <u>2</u> | 0.094 |
| AR-0961 | *Moraxella catarrhalis* | 0.19 | <u>1</u> | <u>2</u> | 0.5 | 0.25 | 0.094 |
| AR-0962 | *Moraxella catarrhalis* | 0.094 | 0.012 | 0.064 | 0.75 | 0.25 | 0.125 |

aReported MICs are the mode of n=3 replicates
bIsolates with reduced susceptibilities are indicated by bold and underlined MIC values (CLSI breakpoints: PEN ≥ 2 µg/mL; CRO ≥ 0.5 µg/n CFX ≥ 0.5 µg/mL; TET ≥ 2 µg/mL; AZI ≥ 2 µg/mL; CIP ≥ 1 µg/mL)

**Table 2.** Genome assembly statistics for sequenced strains.

| Isolate | SRA Read Accession | Genome Size (bp) | GC Content (%) | No. of Contigs | Estimated Coverage (x) | No. of ORFs | No. of tRNAs |
| --- | --- | --- | --- | --- | --- | --- | --- |
| AR-0933 | SAMN15454039 | 2,100,787 | 52.6 | 65 | 438 | 2,059 | 52 |
| AR-0934 | SAMN15454040 | 2,102,184 | 52.7 | 64 | 458 | 2,081 | 49 |
| AR-0935 | SAMN15454041 | 2,215,614 | 52.35 | 54 | 470 | 2,224 | 51 |
| AR-0936 | SAMN15454042 | 2,158,950 | 52.38 | 71 | 380 | 2,117 | 50 |
| AR-0937 | SAMN15454043 | 2,138,399 | 52.53 | 69 | 498 | 2,102 | 49 |
| AR-0938 | SAMN15454044 | 2,152,496 | 52.5 | 71 | 491 | 2,124 | 49 |
| AR-0943 | SAMN15454045 | 2,342,297 | 59.34 | 91 | 421 | 2,151 | 52 |
| AR-0945 | SAMN15454046 | 2,572,594 | 53.81 | 45 | 214 | 2,511 | 55 |
| AR-0946 | SAMN15454047 | 2,145,323 | 52.48 | 53 | 366 | 1,994 | 54 |
| AR-0947 | SAMN15454048 | 2,497,075 | 52.8 | 19 | 247 | 2,302 | 56 |
| AR-0948 | SAMN15454049 | 2,399,311 | 48.41 | 42 | 341 | 2,261 | 52 |
| AR-0949 | SAMN15454050 | 2,938,382 | 50.84 | 98 | 228 | 2,749 | 58 |
| AR-0950 | SAMN15454051 | 1,762,877 | 51.47 | 18 | 344 | 2,184 | 55 |
| AR-0951 | SAMN15454052 | 2,580,111 | 51.1 | 98 | 295 | 2,330 | 54 |
| AR-0952 | SAMN15454053 | 2,446,961 | 51.04 | 66 | 319 | 2,216 | 53 |
| AR-0953 | SAMN15454054 | 2,201,301 | 49.57 | 16 | 921 | 2,096 | 54 |
| AR-0954 | SAMN15454055 | 2,359,025 | 49.13 | 57 | 390 | 2,276 | 54 |
| AR-0955 | SAMN15454056 | 2,188,966 | 49.37 | 16 | 327 | 2,033 | 55 |
| AR-0956 | SAMN15454057 | 2,238,415 | 49.02 | 34 | 397 | 2,106 | 55 |
| AR-0957 | SAMN15454058 | 2,187,919 | 49.37 | 15 | 326 | 2,029 | 55 |
| AR-0958 | SAMN15454059 | 2,268,810 | 49.1 | 30 | 478 | 2,132 | 49 |
| AR-0959 | SAMN15454060 | 2,148,848 | 49.01 | 24 | 382 | 1,962 | 49 |
| AR-0961 | SAMN15454061 | 1,882,811 | 41.81 | 13 | 577 | 1,728 | 44 |
| AR-0962 | SAMN15454062 | 1,843,431 | 41.64 | 12 | 432 | 1,684 | 43 |

Of the typically human-associated commensal *Neisseria* within the panel (n=11), eight displayed reduced susceptibly to at least one of the tested antibiotics (Table 1). Two isolates were resistant to penicillin (*N. bacilliformis* AR Bank # 0943 and *N. subflava* AR Bank # 0954), one was resistant to ceftriaxone (*N. bacilliformis* AR Bank # 0943), two were resistant to cefixime (*N. bacilliformis* AR Bank # 0943 and *N. subflava* AR Bank # 0954), five were resistant to tetracycline (*N. bacilliformis* AR Bank # 0943, *N. cinerea* AR Bank # 0944, *N. subflava* AR Bank # 0954, *N. subflava* AR Bank # 0955, and *N. subflava* AR Bank # 0957), eight were resistant to azithromycin (*N. bacilliformis* AR Bank # 0943, *N. cinerea* AR Bank # 0944, *N. mucosa* AR Bank # 0951, and *N. subflava* AR Bank # 0953 through 0957), and one was resistant to ciprofloxacin (*N. bacilliformis* AR Bank # 0943). Only one isolate was resistant to all test antibiotics, *N. bacilliformis* AR Bank # 0943, which was the only representative of *N. bacilliformis* in the panel. This isolate also had a high MIC to ceftriaxone of 16 μg/mL, with recorded MICs for *N. gonorrhoeae* typically ≤ 1 μg/mL [22]. Two out of the three isolates of the other represented human-associated *Neisseria* (*Moraxella* and *Kingella*) showed reduced susceptibility to azithromycin (*K. denitrificans* AR Bank # 0960), or ceftriaxone and cefixime (*M. catarrhalis* AR Bank # 0961).

For the *Neisseria* typically associated with animals (*N. canis*, *N. macacae*, and *N. weaveri*; n=6), three were resistant to at least one antibiotic (Table 1). *N. macacae* AR Bank # 0949 was resistant to tetracycline, azithromycin, and was just below the reduced susceptibility threshold for penicillin (1.5 μg/mL with the breakpoint at 2 μg/mL); *N. macacae* AR Bank # 0950 was resistant to penicillin, tetracycline, and azithromycin; *N. macacae* AR Bank # 0952 was resistant to tetracycline.

### Genome assemblies and computational analyses

Of the 26 isolates within our panel, we were able to generate genomic libraries for 24 of them. For the remaining isolates, *N. cinerea* AR Bank # 0944 and *K. denitrificans* AR Bank # 0960, we were unable to generate sufficient DNA for sequencing. Genome assemblies ranged from 1.76 to 2.9 Mbp long and contained between 1,684 to 2,749 predicted open reading frames. Coverages ranged from 214x to 921x. All of the sequenced AR Bank isolates clustered with representatives of their own species as assessed via a phylogeny generated from 16S rRNA gene sequences (Figure 1). However, representatives of *N. mucosa* (n=3) and *N. macacae* (n=3) formed a single monophyletic cluster.

**Figure 1.**
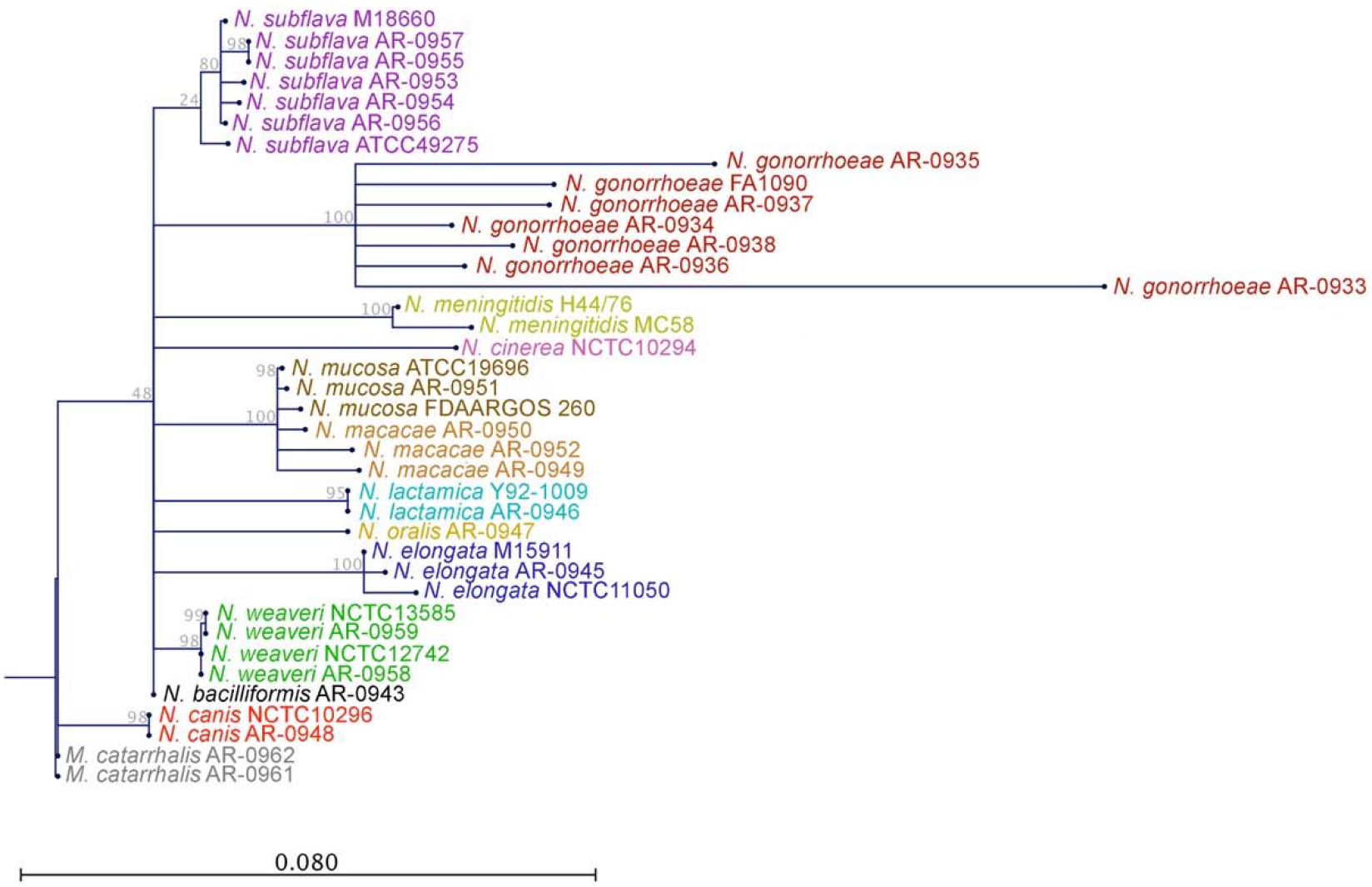
Maximum likelihood phylogenetic tree of 38 *Neisseria* isolates based on the 16S rRNA gene. Species are coded by unique colors. The scale bar represents 0.08 substitutions per nucleotide site. AR Bank # 0961, one of the *Moraxella catarrhalis* in the *Neisseria* species MALDI-TOF Verification panel, was used as an outgroup.

In total, 279 total mutations or haplotypes predicted to encode resistance were found across all sequenced isolates – consisting of 63 unique genetic mechanisms (Figure 2; Supplementary Table 1). These polymorphisms were predicted to encode resistance to 14 different classes of antimicrobial compounds, and included predicted phenotypic resistance to all of the tested drugs within this study.

**Figure 2.**
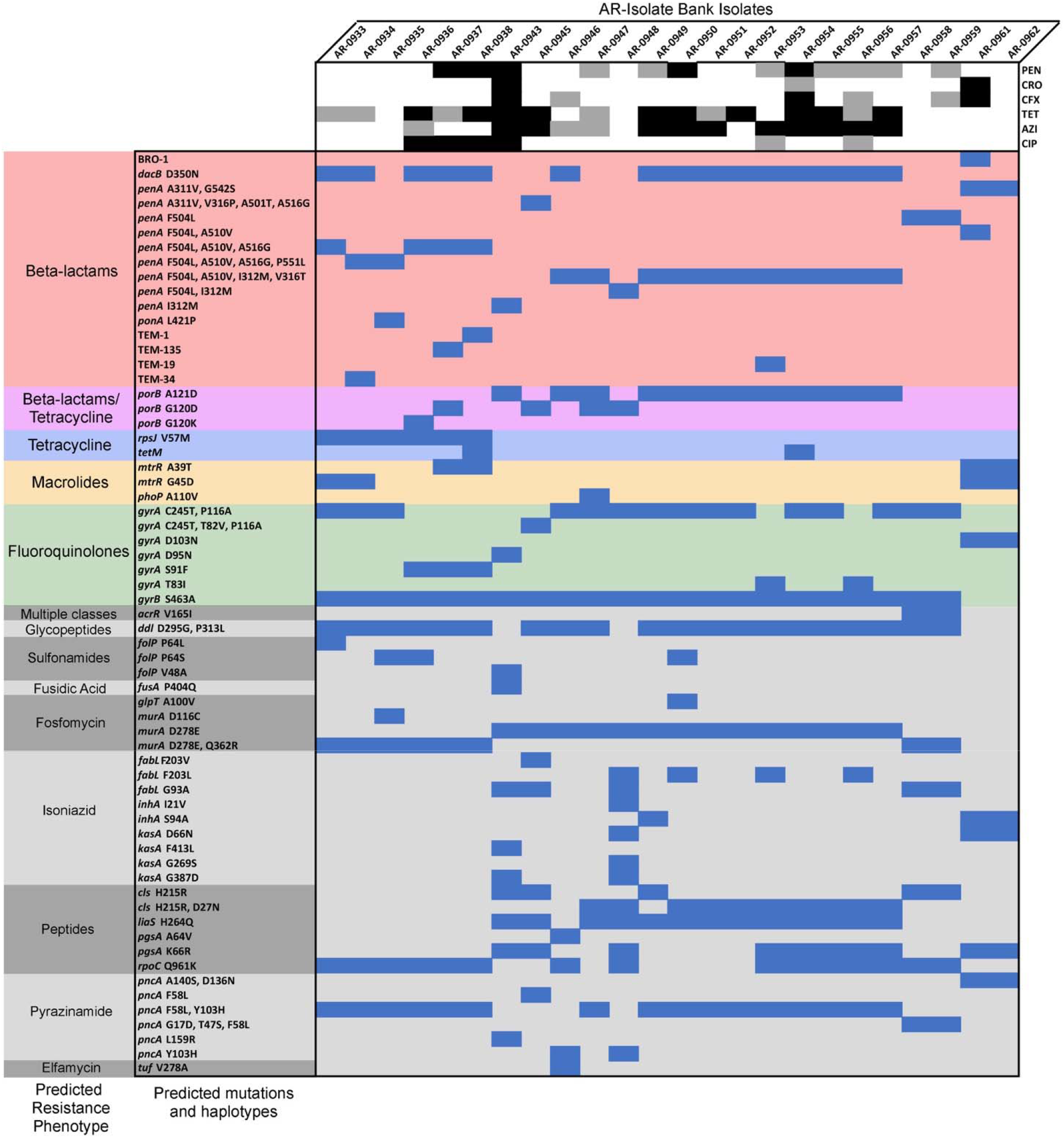
Heatmap showing the distribution of CARD RGI predicted resistance-encoding mutations for all sequenced isolates (n=24). Upper panel, *Neisseria* species’ susceptibility phenotypes as quantified by MIC. Black denotes loss of susceptibility (CLSI standards), grey denotes one dilution below the reduced susceptibility breakpoint, white denotes susceptibility to the antibiotics listed on the right (PEN, penicillin; CRO, ceftriaxone; CFX, cefixime; TET, tetracycline; AZI, azithromycin; and CIP, ciprofloxacin). Lower panel, presence of each recovered mutation is indicated by blue fill. CARD predicted resistance phenotypes for sets of mutations are denoted for beta-lactam resistance (red shading), beta-lactam and tetracycline resistance (purple), tetracycline (blue), macrolides (orange), fluoroquinolones (green), and resistance to other antibiotics (grey).

### Quinolones reduced susceptibility

Resistance to quinolones like ciprofloxacin in gonococci is mediated by mutations that reduce binding affinity to the primary drug target, DNA gyrase subunit A (GyrA) – a type II topoisomerase that negatively supercoils ds-DNA. Specifically, these mutations include either a missense mutation at codon 91 in *gyrA* (S91F) or codon 95 (D95N), which have nearly perfect sensitivity and specificity for the prediction of resistance to this drug in clinical *N. gonorrhoeae* specimens [22,24–27]. In our panel, we documented four isolates with reduced susceptibility to ciprofloxacin (*N. gonorrhoeae* AR Bank # 0936 through 0938, and *N. bacilliformis* AR Bank # 0943), and unsurprisingly they all had either the GyrA S91F or D95N substitutions (Figure 2). However, interestingly, we also find two *N. subflava* isolates (AR Bank # 0953 and AR Bank # 0956) with ciprofloxacin MICs less than one dilution below the reduced susceptibility threshold (Table 1) which did not harbor these mutations. Instead, they had a missense mutation at codon 83 in *gyrA* (T83I), which is located in a quinolone resistance-determining region (QRDR), and has been shown to moderately increase quinolone resistance in *Pseudomonas aeruginosa* [28].

### Macrolide resistance

Macrolide class antibiotics obstruct protein synthesis by binding to the 50S ribosomal subunit, and within *N. gonorrhoeae*, several types of mutations have been documented to be involved in resistance. These mutations include substitutions in the 23S rRNA azithromycin binding sites (C2611T and A2059G) [29,30], the presence of mosaic *multiple transferable resistance* (*mtr*) efflux pump alleles acquired from commensal *Neisseria* species [10,20,22,31], mutations that increase the expression of *mtrCDE* [32–34], mutations in *rplD* [22], *rplV* tandem duplications [22], and variants of the rRNA methylase genes *ermC* and *ermB* [35]. However, we find no isolates in this panel with any of these mutations above the CLSI reduced susceptibility breakpoint (Figure 2). We do find four *N. gonorrhoeae* isolates with missense mutations at codons 39 (A39T) and 45 (G45D) of the repressor of the Mtr efflux pump (MtrR) – which have been shown to enhance *mtrCDE* expression by introducing radical amino acid substitutions in the DNA-binding motif of the repressor, and ablating its promoter-binding function [32,36]. However, these isolates did not have elevated MICs to azithromycin.

### Reduced susceptibility to tetracycline

Tetracycline is a broad-spectrum polyketide antibiotic that binds the bacterial 30S ribosomal subunit and blocks incoming aminoacyl tRNAs from entering the ribosome acceptor site. Here, we find two isolates *N. gonorrhoeae* AR Bank # 0938 and *N. subflava* AR Bank # 0954 with high level tetracycline resistance (≥ 32 μg/mL; Table 1). High level tetracycline resistance in *Neisseria* has been demonstrated to be a direct result of inheritance of a Class M tetracycline resistance determinant (*tetM*), which after binding to the ribosome triggers the release of tetracycline due to its resemblance to elongation factor G (EF-G). Both *N. gonorrhoeae* AR Bank # 0938 and *N. subflava* AR Bank # 0954 had a *tetM* gene present (Figure 2), further supporting prior literature suggesting that *tetM* circulates within commensal *Neisseria* communities [37] in addition to gonococcal populations [38,39]. Lower level tetracycline resistance has been shown to be mediated by mutations that decrease the influx of tetracycline through the porin [40,41], mutations in *mtr* that increase pump expression [41–43], or structure modifying mutations in the ribosome [44]. Several isolates had mutations in the porin (PorB G120K, A121D, or A121N) and/or a mutations in the ribosomal protein RpsJ (V57M) which have previously been associated with reduced susceptibility to tetracycline in previous studies [40,44], however inheritance of these mutations were not perfectly correlated with reduced susceptibility (Figure 2).

### Resistance to β-lactams

Penicillins (penicillin G) and cephalosporins (ceftriaxone and cefixime) are β-lactam antibiotics which inhibit cell wall biosynthesis by binding the transpeptidase enzymes (penicillin-binding proteins [PBPs]) that form the peptidoglycan cross-links in the bacterial cell wall. High level penicillin resistance in gonococci is typically mediated by the presence of a TEM-1-type β-lactamase encoded by the *bla*_*TEM-1*_ gene, which acts through degradation of the four-atom β-lactam ring of penicillin [45]. Of the isolates within this panel with penicillin MICs ≥ 32 μg/mL, two out of three had a TEM-type β-lactamase present. Lower level penicillin resistance can be modulated through multiple mutations including those in *penA* [41,46], *mtr* and its regulatory components [41–43], *porB* [40,41], *ponA* [47], and *pilQ* [47,48] – which often contribute additively to one another [47]. Notably within the commensals in this panel we find at least one of these mutations present in all isolates (i.e., *porB* G120K, A121D, or A121N; *ponA* L421P; or several mutations in *penA* [Figure 2]), suggesting their widespread availability for horizontal exchange.

In contrast to the multiple mutations that give rise to penicillin resistance, reduced susceptibility to cephalosporins in *Neisseria* is most frequently due to mutations in the PBP targets of the drug which decrease their acylation rate. Commensal *Neisseria spps.* are proven sources of alleles conferring cephalosporin resistance for *N. gonorrhoeae*, and within this panel we find *penA* alleles with several amino acid substitutions that have been proven or associated with increased resistance including: A311V, I312M, V316T, V316P, A501T, G542S, and P551L [21,49–51] (Figure 2). Finally, we found a BRO-1 β-lactamase in one of the *Moraxella catarrhalis* isolates (AR Bank # 0961) within the panel which also was resistant to both cefixime and ceftriaxone (Table 1, Figure 2). The BRO-1 β-lactamase has previously been associated with resistance to β-lactams, and is present in over 90% of all *M. catarrhalis* isolates [52,53], though it is unclear if this β-lactamase can be transferred to the other *Neisseria*.

## Discussion

The significance of the commensal *Neisseria* as reservoirs of antibiotic resistance for gonococci has been repeatedly demonstrated [10,21,22,49–51,54], which emphasizes the importance of characterizing the mutations encoding reduced susceptibility across the entirety of this species consortium. Our results offer an initial exploration into the *Neisseria* resistome, and clearly demonstrate resistance-encoding mutations that are known in gonococci (*gyrA* (S91F), *tetM*, and TEM-type β-lactamases) also circulate in commensal communities, suggesting their widespread availability for horizontal transfer. However, it is also clear that we have not captured all mutations involved in producing reduced susceptibility across this panel. This is evidence by isolates with the same reported resistance haplotypes displaying variation in MIC. For example, *N. subflava* AR Bank # 0954 and 0955 had MICs to penicillin of 3 and 1 μg/mL respectively, despite having the same mutations in *penA* and *porB* (Table 1 and Figure 2). Similarly, *N. gonorrhoeae* AR Bank # 0936 and 0937 isolates had ciprofloxacin MICs of 32 and 6 μg/mL respectively, yet harbored the same *gyrA* (S91F) substitution (Table 1 and Figure 2). Furthermore, there are cases of unexplained resistance for which no putative resistance mutations had been identified – especially for the drug azithromycin. However, this is not necessarily surprising for macrolide antibiotics, as much of the genetic basis of resistance remains unclear in gonococci [22], suggesting that many mutational steps are required and that there may be many different paths to reduced susceptibility. Ultimately, these discrepancies between phenotype and genotype point to the main limitations of our approach, which include: the dependence on a database that contains the full universe of polymorphisms within a given species or genus that may give rise to resistance, sufficient sequence homology between database entries and query sequences for alignment, and knowledge of epistatic modulators of resistance that may not directly impact MIC on their own but only in combination with other mutations.

Though lab-based efforts will be needed to confirm the causality of our CARD-nominated mutations in reduced susceptibility in addition to the elucidation of the genetic underpinnings that to contribute to the unexplained MIC variance, our combined approaches employing both experimental quantification of MIC data coupled with genomic sequencing is a key first step in exploring the possible resistance mechanisms harbored by commensal populations. Ultimately, these types of studies will provide the foundation for prospective surveillance of novel resistance determinants that may be rapidly acquired by pathogens of critical importance across a wide range of genera.

## Materials and methods

### Bacterial strains and culture conditions

All isolates were cultured on GCB agar medium (Becton Dickinson Co., Franklin Lakes, NJ) supplemented with 1% Kellogg’s solution (GCB-K plates) [55] at 37°C in a 5% CO_2_ incubator. Stocks for all bacteria were stored at −80°C in trypticase soy broth containing 50% glycerol.

### Minimum inhibitory concentration testing

Antimicrobial susceptibility testing was conducted using Etest strips on GCB-K plates, according to the manufacturer specifications (bioMérieux, Durham, NC) – which have been shown to have comparable MIC values to the agar dilution method, with the exception of cefixime for which Etests systemically report lower MICs [56]. In brief, cells from overnight plates were suspended in trypticase soy broth to a 0.5 McFarland standard and inoculated onto new GCB-K plates. Etest strips were subsequently placed on the surface of the inoculated plates. Following 18-24 hours of incubation at 37°C in a 5% CO_2_ incubator, MICs were determined by reading the lowest concentration that inhibited growth, and reduced susceptibility was determined using CLSI guidelines (CLSI breakpoints: PEN ≥ 2 μg/mL; CRO ≥ 0.5 μg/mL; CFX ≥ 0.5 μg/mL; TET ≥ 2 μg/mL; AZI ≥ 2 μg/mL; CIP ≥ 1 μg/mL) [57]. MICs were read by at least two independent researchers, and the mode of three tests was reported.

### Library preparation and genomic sequencing

Genomic DNA was isolated by lysing growth from overnight plates in TE buffer (10 mM Tris [pH 8.0], 10 mM EDTA) with 0.5 mg/mL lysozyme and 3 mg/mL proteinase K (Sigma-Aldrich Corp., St. Louis, MO). To obtain sufficient DNA from AR BANK # 0943 and AR BANK # 0958, which were difficult to swab off of GCB-K plates, we instead inoculated liquid GCB broth media (7.5 g protease peptone #3, 0.5 g soluble starch, 2 g dibasic K_2_HPO_4_, 0.5 g monobasic KH_2_PO_4_, 2.5 g NaCl, ddH_2_O to 500mL; Becton Dickinson) supplemented with 1% Kellogg’s solution and incubated overnight at 37°C. After 24 hours, cultures were centrifuged for 10 minutes at 14,000 rpm. The supernatant was discarded and the same method as described above was used to isolate DNA.

DNA was purified using the PureLink Genomic DNA Mini kit (Thermo Fisher Corp., Waltham, MA), treated with RNase A, and stored in water. Sequencing libraries were prepared using the Nextera XT kit as per the manufacturer’s instructions (Illumina Corp., San Diego, CA). Samples were uniquely dual-indexed and pooled (n = 17-18 libraries per pool), and sequenced using a V3 600 cycle cartridge (2×300bp) on an Illumina MiSeq platform at the Rochester Institute of Technology Genomics Core.

### Genome assembly and bioinformatic analyses

Sequencing quality of each paired-end read library was assessed using FastQC v0.11.9 [58]. Trimmomatic v0.39 [59] was used to trim adapter sequences, and remove bases with phred quality score < 15 over a 4 bp sliding window. Reads < 36 bp long, or those missing a mate, were also removed from subsequent analysis. Trimmed reads were assembled using SPAdes v.3.7.0 [60], and the resultant *de novo* assemblies were evaluated using the Quality Assessment Tool for Genome Assemblies (QUAST) v.4.1 [61]. Prokka v.1.11 [62] was used to annotate assemblies. Resistance-encoding mutations were predicted using the Comprehensive Antibiotic Resistance Database (CARD) Resistance Gene Identifier (RGI) v5.1.0 [63]. Gene presence or absence was not considered, unless *tetM* or TEM beta lactamases were recorded, which are most often harbored on plasmids [64].

To assess evolutionary relationships between isolates, we used the annotated 16s rRNA gene in FA1090 (AE004969.1) as a reference, and subset the homologous region in each assembly with blastn. We then used CLC Main Workbench v20.0.4 [65] to align sequences and reconstruct a maximum likelihood phylogeny using 100 bootstrap replicates and the Jukes Cantor substitution model.

## Data availability

All genomic read libraries generated in this study are available from the NCBI SRA database (accession numbers SAMN15454039-SAMN15454062). Scripts for analyses can be found at https://github.com/wadsworthlab.

## Acknowledgements

The authors would like to thank the Thomas H. Gosnell School of Life Sciences (GSoLS) and the College of Science (COS) at the Rochester Institute of Technology (RIT) for ongoing support of this work. We would also like to thank the CDC & FDA’s Antibiotic Resistance (AR) Isolate Bank for generously providing the isolates used in this study. A.O.H acknowledges support from the National Institutes of Health (NIH) award (R15GM120653). The authors thank Maira Goytia (Spelman College) for providing feedback on an earlier draft of this manuscript.

## Author contributions

C.B.W. conceptualized and provided the resources for the study. C.B.W., M.F., J.R., and N.W. performed experiments and data analyses. C.B.W. and A.O.H. assisted with data interpretation and preparation of the manuscript. All authors contributed to and approved the final manuscript. This study was supported by the Thomas H. Gosnell School of Life Sciences and the College of Science at RIT.

## References

1. U.S. Centers for Disease Control and Prevention Antibiotic Resistance Threats in the United States, 2019. 2019, 1–148.

2. Walsh, F.; Duffy, B. The culturable soil antibiotic resistome: A community of multi-drug resistant bacteria. PLoS ONE 2013, 8, e65567.

3. Forsberg, K. J.; Reyes, A.; Wang, B.; Selleck, E. M.; Sommer, M. O. A.; Dantas, G. The shared antibiotic resistome of soil bacteria and human pathogens. Science 2012, 337, 1107–1111.

4. D’Costa, V. M.; McGrann, K. M.; Hughes, D. W.; Wright, G. D. Sampling the antibiotic resistome. Science 2006, 311, 374–377.

5. Bezanson, G. S.; Khakhria, R.; Bollegraaf, E. Nosocomial outbreak caused by antibiotic-resistant strain of *Salmonella typhimurium* acquired from dairy cattle. Can. Med. Assoc. J. 1983, 128, 426–427.

6. Spika, J. S.; Waterman, S. H.; Hoo, G. W.; St Louis, M. E.; Pacer, R. E.; James, S. M.; Bissett, M. L.; Mayer, L. W.; Chiu, J. Y.; Hall, B. Chloramphenicol-resistant *Salmonella newport* traced through hamburger to dairy farms. N. Engl. J. Med. 1987, 316, 565–570.

7. Levy, S. B.; FitzGerald, G. B.; Macone, A. B. Changes in intestinal flora of farm personnel after introduction of a tetracycline-supplemented feed on a farm. N. Engl. J. Med. 1976, 295, 583–588.

8. Rashid, H.; Rahman, M. Possible transfer of plasmid mediated third generation cephalosporin resistance between *Escherichia coli* and *Shigella sonnei* in the human gut. Infect. Gent. Evol. 2015, 30, 15–18.

9. Knudsen, P. K.; Gammelsrud, K. W.; Alfsnes, K.; Steinbakk, M.; Abrahamsen, T. G.; ller, F. M. X.; Bohlin, J. Transfer of a blaCTX-M-1-carrying plasmid between different *Escherichia coli* strains within the human gut explored by whole genome sequencing analyses. Sci. Rep. 2017, 1–10.

10. Wadsworth, C. B.; Arnold, B. J.; Sater, M. R. A.; Grad, Y. H. Azithromycin resistance through interspecific acquisition of an epistasis-dependent efflux pump component and transcriptional regulator in *Neisseria gonorrhoeae*. mBio 2018, 9, 1–17.

11. Blair, J. M. A.; Webber, M. A.; Baylay, A. J.; Ogbolu, D. O.; Piddock, L. J. V. Molecular mechanisms of antibiotic resistance. Nat. Rev. Microbiol. 2014, 13, 42–51.

12. Eyre, D. W.; Sanderson, N. D.; Lord, E.; Regisford-Reimmer, N.; Chau, K.; Barker, L.; Morgan, M.; Newnham, R.; Golparian, D.; Unemo, M.; Crook, D. W.; Peto, T. E.; Hughes, G.; Cole, M. J.; Fifer, H.; Edwards, A.; Andersson, M. I. Gonorrhoea treatment failure caused by a *Neisseria gonorrhoeae* strain with combined ceftriaxone and high-level azithromycin resistance, England, February 2018. Eurosurveillance 2018, 23, 364–6.

13. Jennison, A. V.; Whiley, D.; Lahra, M. M.; Graham, R. M.; Cole, M. J.; Hughes, G.; Fifer, H.; Andersson, M.; Edwards, A.; Eyre, D. Genetic relatedness of ceftriaxone-resistant and high-level azithromycin resistant *Neisseria gonorrhoeae* cases, United Kingdom and Australia, February to April 2018. Eurosurveillance 2019, 24, 717–4

14. Corander, J.; Connor, T. R.; O’Dwyer, C. A.; Kroll, J. S.; Hanage, W. P. Population structure in the *Neisseria*, and the biological significance of fuzzy species. J. R. Soc. Interface 2011, 9, 1208–1215.

15. Halter, R.; Pohlner, J.; Meyer, T. F. Mosaic-like organization of *igA protease* genes in *Neisseria gonorrhoeae* generated by horizontal genetic exchange *in vivo*. EMBO J. 1989, 8, 2737–2744.

16. Feavers, I. M.; Heath, A. B.; Bygraves, J. A.; Maiden, M. C. Role of horizontal genetic exchange in the antigenic variation of the class 1 outer membrane protein of *Neisseria meningitidis*. Mol. Microbiol. 1992, 6, 489–495.

17. Smith, J. M.; Smith, N. H.; O’Rourke, M.; Spratt, B. G. How clonal are bacteria? P. Natl. A. Sci. 1993, 90, 4384–4388.

18. Zhou, J.; Spratt, B. G. Sequence diversity within the *argF*, *fbp* and *recA* genes of natural isolates of *Neisseria meningitidis*: Interspecies recombination within the *argF* gene. Mol. Microbiol. 1992, 6, 2135–2146.

19. Arnold, B.; Sohail, M.; Wadsworth, C.; Corander, J.; Hanage, W. P.; Sunyaev, S.; Grad, Y. H. Fine-Scale haplotype structure reveals strong signatures of positive selection in a recombining bacterial pathogen. Mol. Biol. Evol. 2020, 37, 417–428.

20. Rouquette-Loughlin, C. E.; Reimche, J. L.; Balthazar, J. T.; Dhulipala, V.; Gernert, K. M.; Kersh, E. N.; Pham, C. D.; Pettus, K.; Abrams, A. J.; Trees, D. L.; St Cyr, S.; Shafer, W. M. Mechanistic basis for decreased antimicrobial susceptibility in a clinical isolate of *Neisseria gonorrhoeae* oossessing a mosaic-Like *mtr* efflux pump locus. mBio 2018, 9, 587.

21. Spratt, B. G.; Bowler, L. D.; Zhang, Q. Y.; Zhou, J.; Smith, J. M. Role of interspecies transfer of chromosomal genes in the evolution of penicillin resistance in pathogenic and commensal *Neisseria* species. J. Mol. Evol. 1992, 34, 115–125.

22. Grad, Y. H.; Harris, S. R.; Kirkcaldy, R. D.; Green, A. G.; Marks, D. S.; Bentley, S. D.; Trees, D.; Lipsitch, M. Genomic epidemiology of gonococcal resistance to extended-spectrum cephalosporins, macrolides, and fluoroquinolones in the United States, 2000-2013. J. Infect. Dis. 2016, 214, 1579–1587.

23. CDC & FDA Antibiotic Resistance Isolate Bank. Atlanta (GA): CDC. July 22, 2020.

24. Lindbäck, E.; Rahman, M.; Jalal, S.; Wretlind, B. Mutations in *gyrA, gyrB, parC*, and *parE* in quinolone-resistant strains of *Neisseria gonorrhoeae*. APMIS 2002, 110, 651–657.

25. Peterson, S. W.; Martin, I.; Demczuk, W.; Bharat, A.; Hoang, L.; Wylie, J.; Allen, V.; Lefebvre, B.; Tyrrell, G.; Horsman, G.; Haldane, D.; Garceau, R.; Wong, T.; Mulvey, M. R. Molecular assay for detection of ciprofloxacin resistance in *Neisseria gonorrhoeae* isolates from cultures and clinical nucleic acid amplification test specimens. J. Clin. Microbiol. 2015, 53, 3606–3608.

26. Hemarajata, P.; Yang, S.; Soge, O. O.; Humphries, R. M.; Klausner, J. D. Performance and verification of a real-time pcr assay targeting the *gyrA* gene for prediction of ciprofloxacin resistance in *Neisseria gonorrhoeae*. J. Clin. Microbiol. 2016, 54, 805–808.

27. Ebeyan, S.; Windsor, M.; Bordin, A.; Mhango, L.; Erskine, S.; Mokany, E.; Tan, L. Y.; Whiley, D.; GRAND2 Study Investigators; Guy, R.; Wand, H.; Donovan, B.; Bell, S.; Kaldor, J.; Kong, M.; Conway, D.; Lafferty, L.; Ward, J.; Lahra, M.; Chen, M.; Ryder, N.; Lewis, D.; Paterson, D.; Trembizki, E.; Baird, R.; Fairley, C.; Gunathilake, M.; Howden, B.; Klausner, J.; Nimmo, G.; Russell, D.; Menon, A.; Palmer, C.; McNulty, A.; Templeton, D.; Cunningham, P.; van Hal, S.; Givney, R. Evaluation of the ResistancePlus GC (beta) assay: a commercial diagnostic test for the direct detection of ciprofloxacin susceptibility or resistance in *Neisseria gonorrhoeae*. J. Antimicrob. Chemoth. 2019, 74, 1820–1824.

28. Bruchmann, S.; Dötsch, A.; Nouri, B.; Chaberny, I. F.; Häussler, S. Quantitative contributions of target alteration and decreased drug accumulation to *Pseudomonas aeruginosa* fluoroquinolone resistance. Antimicrob. Agents Ch. 2013, 57, 1361–1368.

29. Ng, L. K.; Martin, I.; Liu, G.; Bryden, L. Mutation in 23S rRNA associated with macrolide resistance in *Neisseria gonorrhoeae*. Antimicrob. Agents Ch. 2002, 46, 3020–3025.

30. Chisholm, S. A.; Dave, J.; Ison, C. A. High-Level azithromycin resistance occurs in *Neisseria gonorrhoeae* as a result of a single point mutation in the 23S rRNA genes. Antimicrob. Agents Ch. 2010, 54, 3812–3816.

31. Trembizki, E.; Doyle, C.; Jennison, A.; Smith, H.; Bates, J.; Lahra, M.; Whiley, D. A *Neisseria gonorrhoeae* strain with a meningococcal *mtrR* sequence. J. Med. Microbiol. 2014, 63, 1113–1115.

32. Zarantonelli, L.; Borthagaray, G.; Lee, E. H.; Veal, W.; Shafer, W. M. Decreased susceptibility to azithromycin and erythromycin mediated by a novel *mtr(R)* promoter mutation in *Neisseria gonorrhoeae*. J. Antimicrob. Chemoth. 2001, 47, 651–654.

33. Ohneck, E. A.; Zalucki, Y. M.; Johnson, P. J. T.; Dhulipala, V.; Golparian, D.; Unemo, M.; Jerse, A. E.; Shafer, W. M. A novel mechanism of high-level, broad-spectrum antibiotic resistance caused by a single base pair change in *Neisseria gonorrhoeae*. mBio 2011, 2, e00187–11–e00187–11.

34. Hagman, K. E.; Pan, W.; Spratt, B. G.; Balthazar, J. T.; Judd, R. C.; Shafer, W. M. Resistance of *Neisseria gonorrhoeae* to antimicrobial hydrophobic agents is modulated by the *mtrRCDE* efflux system. Microbiology 1995, 141, 611–622.

35. Demczuk, W.; Martin, I.; Peterson, S.; Bharat, A.; Van Domselaar, G.; Graham, M.; Lefebvre, B.; Allen, V.; Hoang, L.; Tyrrell, G.; Horsman, G.; Wylie, J.; Haldane, D.; Archibald, C.; Wong, T.; Unemo, M.; Mulvey, M. R. Genomic epidemiology and molecular resistance mechanisms of azithromycin-resistant *Neisseria gonorrhoeae* in Canada from 1997 to 2014. J. Clin. Microbiol. 2016, 54, 1304–1313.

36. Shafer, W. M.; Balthazar, J. T.; Hagman, K. E.; Morse, S. A. Missense mutations that alter the DNA-binding domain of the MtrR protein occur frequently in rectal isolates of *Neisseria gonorrhoeae* that are resistant to faecal lipids. Microbiology 1995, 141 (Pt 4), 907–911.

37. Knapp, J. S.; Johnson, S. R.; Zenilman, J. M.; Roberts, M. C.; Morse, S. A. High-level tetracycline resistance resulting from TetM in strains of *Neisseria spp., Kingella denitrificans*, and *Eikenella corrodens*. Antimicrob. Agents Ch. 1988, 32, 765–767.

38. Morse, S. A.; Johnson, S. R.; Biddle, J. W.; Roberts, M. C. High-level tetracycline resistance in *Neisseria gonorrhoeae* is result of acquisition of streptococcal *tetM* determinant. Antimicrob. Agents Ch. 1986, 30, 664–670.

39. Gascoyne, D. M.; Heritage, J.; Hawkey, P. M.; Turner, A.; van Klingeren, B. Molecular evolution of tetracycline-resistance plasmids carrying TetM found in *Neisseria gonorrhoeae* from different countries. J. Antimicrob. Chemoth. 1991, 28, 173–183.

40. Gill, M. J.; Simjee, S.; Al-Hattawi, K.; Robertson, B. D.; Easmon, C. S. F.; Ison, C. A. Gonococcal resistance to β-lactams and tetracycline involves mutation in loop 3 of the porin encoded at the *penB* locus. Antimicrob. Agents Ch. 1998, 42, 2799–2803.

41. Sparling, P. F.; Sarubbi, F. A.; Blackman, E. Inheritance of low-level resistance to penicillin, tetracycline, and chloramphenicol in *Neisseria gonorrhoeae*. J, Bacteriol. 1975, 124, 740–749.

42. Pan, W.; Spratt, B. G. Regulation of the permeability of the gonococcal cell envelope by the *mtr* system. Mol. Microbiol. 1994, 11, 769–775.

43. Veal, W. L.; Nicholas, R. A.; Shafer, W. M. Overexpression of the MtrC-MtrD-MtrE efflux pump due to an *mtrR* mutation is required for chromosomally mediated penicillin resistance in *Neisseria gonorrhoeae*. J, Bacteriol. 2002, 184, 5619–5624.

44. Hu, M.; Nandi, S.; Davies, C.; Nicholas, R. A. High-Level chromosomally mediated tetracycline resistance in *Neisseria gonorrhoeae* results from a point mutation in the *rpsJ* gene encoding ribosomal protein S10 in combination with the *mtrR* and *penB* resistance determinants. Antimicrob. Agents Ch. 2005, 49, 4327–4334.

45. Ashford, W.; Golash, R.; Hemming, V. Penicilunase-producing *Neisseria gonorrhoeae*. Lancet 2020, 308, 657–658.

46. Dowson, C. G.; Jephcott, A. E.; Gough, K. R.; Spratt, B. G. Penicillin-binding protein 2 genes of non-beta-lactamase-producing, penicillin-resistant strains of *Neisseria gonorrhoeae*. Mol. Microbiol. 1989, 3, 35–41.

47. Ropp, P. A.; Hu, M.; Olesky, M.; Nicholas, R. A. Mutations in *ponA*, the gene encoding Penicillin-Binding Protein 1, and a novel locus, *penC*, are required for high-level chromosomally mediated penicillin resistance in *Neisseria gonorrhoeae*. Antimicrob. Agents Ch. 2002, 46, 769–777.

48. Zhao, S.; Tobiason, D. M.; Hu, M.; Seifert, H. S.; Nicholas, R. A. The *penC* mutation conferring antibiotic resistance in *Neisseria gonorrhoeae* arises from a mutation in the PilQ secretin that interferes with multimer stability. Mol. Microbiol. 2005, 57, 1238–1251.

49. Ohnishi, M.; Golparian, D.; Shimuta, K.; Saika, T.; Hoshina, S.; Iwasaku, K.; Nakayama, S. I.; Kitawaki, J.; Unemo, M. Is *Neisseria gonorrhoeae* initiating a future era of untreatable gonorrhea?: Detailed characterization of the first strain with high-level resistance to ceftriaxone. Antimicrob. Agents Ch. 2011, 55, 3538–3545.

50. Ameyama, S.; Onodera, S.; Takahata, M.; Minami, S.; Maki, N.; Endo, K.; Goto, H.; Suzuki, H.; Oishi, Y. Mosaic-Like structure of Penicillin-Binding Protein 2 gene (*penA*) in clinical isolates of *Neisseria gonorrhoeae* with reduced susceptibility to cefixime. Antimicrob. Agents Ch. 2002, 46, 3744–3749.

51. Ito, M.; Deguchi, T.; Mizutani, K.-S.; Yasuda, M.; Yokoi, S.; Ito, S.-I.; Takahashi, Y.; Ishihara, S.; Kawamura, Y.; Ezaki, T. Emergence and spread of *Neisseria gonorrhoeae* clinical isolates harboring mosaic-like structure of Penicillin-Binding Protein 2 in Central Japan. Antimicrob. Agents Ch. 2005, 49, 137–143.

52. Fung, C. P.; Yeo, S. F.; Livermore, D. M. Susceptibility of *Moraxella catarrhalis* isolates to beta-lactam antibiotics in relation to beta-lactamase pattern. J. Antimicrob. Chemoth. 1994, 33, 215–222.

53. Khan, M. A.; Northwood, J. B.; Levy, F.; Verhaegh, S. J. C.; Farrell, D. J.; Van Belkum, A.; Hays, J. P. BRO β-lactamase and antibiotic resistances in a global cross-sectional study of *Moraxella catarrhalis* from children and adults. J. Antimicrob. Chemoth. 2009, 65, 91–97.

54. Rouquette-Loughlin, C.; Reimche, J. L.; Balthazar, J. T.; Dhulipala, V.; Gernert, K.; Kersh, E.; Pham, C.; Pettus, K.; Abrams, A. J.; Trees, D. L.; St Cyr, S.; Shafer, W. M. Mechanistic basis for decreased antimicrobial susceptibility in a clinical isolate of *Neisseria gonorrhoeae* possessing a mosaic-like *mtr* efflux pump locus. mBio 2018, 9(6), e02281–18.

55. Kellogg, D. S.; Peacock, W. L.; Deacon, W. E.; Brown, l.; Pirkle, D. *Neisseria gonorrhoeae*. I. Virulence genetically linked to clonal variation. J. Bacteriol. 1963, 85, 1274–1279.

56. Papp, J. R.; Rowlinson, M.-C.; O’Connor, N. P.; Wholehan, J.; Razeq, J. H.; Glennen, A.; Ware, D.; Iwen, P. C.; Lee, L. V.; Hagan, C. Accuracy and reproducibility of the Etest to detect drug-resistant *Neisseria gonorrhoeae* to contemporary treatment. J. Med. Microbiol. 2018, 67, 68–73.

57. Clinical Laboratory Standards Institute. 2019. Performance standards for antimicrobial susceptibility testing, 29th ed. CLSI supplement M100. Clinical and Laboratory Standards Institute, Wayne, PA.

58. Andrews, S. FASTQC: A quality control tool for high throughput sequence data. 2010, 1–7.

59. Bolger, A. M.; Lohse, M.; Usadel, B. Trimmomatic: a flexible trimmer for Illumina sequence data. Bioinformatics 2014, 30, 2114–2120.

60. Bankevich, A.; Nurk, S.; Antipov, D.; Gurevich, A. A.; Dvorkin, M.; Kulikov, A. S.; Lesin, V. M.; Nikolenko, S. I.; Pham, S.; Prjibelski, A. D.; Pyshkin, A. V.; Sirotkin, A. V.; Vyahhi, N.; Tesler, G.; Alekseyev, M. A.; Pevzner, P. A. SPAdes: A New genome assembly algorithm and its applications to single-cell sequencing. J. Comput. Biol. 2012, 19, 455–477.

61. Gurevich, A.; Saveliev, V.; Vyahhi, N.; Tesler, G. QUAST: Quality assessment tool for genome assemblies. Bioinformatics 2013, 29, 1072–1075.

60. Seemann, T. Prokka: rapid prokaryotic genome annotation. Bioinformatics 2014, 30, 2068–2069.

62. Alcock, B. P.; Raphenya, A. R.; Lau, T. T. Y.; Tsang, K. K.; Bouchard, M.; Edalatmand, A.; Huynh, W.; Nguyen, A.-L. V.; Cheng, A. A.; Liu, S.; Min, S. Y.; Miroshnichenko, A.; Tran, H.-K.; Werfalli, R. E.; Nasir, J. A.; Oloni, M.; Speicher, D. J.; Florescu, A.; Singh, B.; Faltyn, M.; Hernandez-Koutoucheva, A.; Sharma, A. N.; Bordeleau, E.; Pawlowski, A. C.; Zubyk, H. L.; Dooley, D.; Griffiths, E.; Maguire, F.; Winsor, G. L.; Beiko, R. G.; Brinkman, F. S. L.; Hsiao, W. W. L.; Domselaar, G. V.; McArthur, A. G. CARD 2020: Antibiotic resistome surveillance with the comprehensive antibiotic resistance database. Nucleic Acids Res. 2019, 10, 226–9.

63. Roberts, M. C. Plasmids of *Neisseria gonorrhoeae* and other *Neisseria* species. Clin. Microbiol. Rev. 1989, 2 Suppl, S18–23.

64. CLC Genomics Workbench 20.0 (https://digitalinsights.qiagen.com); 2020.

